# Ribosome profiling reveals distinct translational programs underlying Arabidopsis seed dormancy and germination

**DOI:** 10.64898/2026.01.08.696037

**Authors:** Maria Victoria Gomez Roldan, Elodie Layat, Julia Bailey-Serres, Jérémie Bazin, Christophe Bailly

**Affiliations:** Sorbonne Université, CNRS, Inserm, Institut de Biologie Paris-Seine, IBPS, F-75005 Paris, France; Center for Plant Cell Biology, Department of Botany and Plant Sciences, University of California, Riverside, CA, USA; IPS2, CNRS, INRAE, Universités Paris-Saclay, Evry and Paris-Cité, F-91190, Gif sur Yvette, France

**Author notes:** Authors for correspondence : Jérémie Bazin; Christophe Bailly.

**Keywords:** *Arabidopsis thaliana*, seed germination, dormancy, translational regulation ribosome profiling (Ribo-seq), upstream open reading frames (uORFs)

## Abstract

Seed dormancy and germination represent a critical developmental transition that determines plant fitness, yet the contribution of translational regulation to this process remains poorly understood. Here, we used genome-wide ribosome profiling (Ribo-seq) combined with RNA sequencing (RNA-seq) to investigate how translational control shapes the transition from dormancy to germination in *Arabidopsis thaliana* seeds. We analyzed dry dormant seeds, stratified non-dormant seeds, and seeds during early imbibition, enabling simultaneous assessment of transcript abundance and ribosome occupancy. Our analyses reveal that dry seeds harbor an unexpectedly organized translational machinery, with ribosomes pre-positioned at start codons and within coding regions of thousands of stored mRNAs, indicating a poised translational state. Dormancy release and early imbibition triggered extensive gene-specific changes in translational efficiency that were largely uncoupled from transcript abundance, highlighting selective translation as a key regulatory layer. Genes involved in ribosome biogenesis, protein folding, and hormone signaling were preferentially translated during dormancy maintenance, whereas germination-promoting factors showed increased ribosome occupancy following stratification. Global ribosome profiling further uncovered dynamic ribosome pausing at stop codons and pronounced modulation of translation initiation during imbibition.We also identified widespread translation of upstream open reading frames (uORFs) and demonstrated that uORF-mediated repression constitutes a major translational checkpoint during seed imbibition. Functional assays confirmed that uORFs from *MARD1* and *PAO4* repress downstream translation *in vivo*. Together, our results establish translational regulation as a central mechanism governing seed dormancy and germination, revealing how ribosome positioning and uORF activity fine-tune protein synthesis to control developmental transitions in response to environmental cues.

## Introduction

Seed germination marks a critical developmental transition in the plant life cycle, determining the timing of seedling establishment and influencing species fitness. This process is governed by a complex interplay of molecular mechanisms that integrate environmental cues, e.g. temperature or light, with endogenous developmental programs, such as seed dormancy. Seed dormancy can be defined as the failure of an intact viable seed to complete germination under apparently favorable conditions (Bewley, 1997). In the model species *Arabidopsis thaliana*, the transcriptional networks regulating seed dormancy and germination have been extensively described. It has been shown that the transition from a dry, quiescent seed to a metabolically active state is associated with major transcriptome reprogramming (Dekkers *et al*., 2013; Narsai *et al*., 2017). Initiation of *de novo* transcription occurs early in germination, within 1 h of the seed imbibing in water, and does not require *de novo* protein synthesis (Kimura & Nambara, 2010). Seed germination in *Arabidopsis* involves two major transcriptional phases separated by testa rupture. The early phase is characterized by extensive transcriptome reprogramming as the seed transitions from a quiescent to a metabolically active state, while a second transcriptional wave accompanies testa rupture and radicle emergence, activating growth- and cell wall-related genes (Dekkers *et al*., 2013). Gene expression studies have revealed the roles of hormonal pathways, particularly abscisic acid (ABA) and gibberellins (GA), and their downstream transcriptional regulators (Bassel *et al*., 2011; Narsai *et al*., 2017). During imbibition, dormant *Arabidopsis* seeds activate a specific transcriptional program that reinforces dormancy rather than promoting germination. ABA-related transcription factors such as *ABI3*, *ABI4*, and *ABI5*, together with *DOG1*, remain highly expressed, maintaining ABA signaling and repressing growth-associated genes. In contrast, non-dormant seeds downregulate these transcripts upon imbibition, enabling the transition toward germination-related transcriptional activity (Nakabayashi *et al*., 2005; Cadman *et al*., 2006; Bentsink *et al*., 2006; Holdsworth *et al*., 2008). However, germination is not solely determined by changes in transcript abundance. Many early cellular events occur without new transcription, relying on the activation of stored mRNAs accumulated during seed maturation. During this step, numerous fully processed mRNAs are synthesized and stored in a translationally inactive state, remaining stable throughout desiccation. Upon imbibition, these pre-existing mRNAs are rapidly translated, enabling early germination to proceed largely independently of *de novo* transcription (Rajjou *et al*., 2004). Later on, during seed imbibition, it has been shown that that mRNA abundance does not directly predict translation activity (Basbouss-Serhal *et al*., 2015). A substantial proportion of transcripts enriched in nondormant and dormant seeds were regulated at the translational rather than transcriptional level. These findings suggest that germination and dormancy are tightly controlled by selective translation.

Translational control, i.e. the regulation at the level of mRNA recruitment to ribosomes, has thus emerged as a pivotal yet poorly explored mechanism in seed biology. Dry quiescent seeds store thousands of mRNAs in a translationally repressed state (so called stored mRNA or long-lived mRNA), and their selective recruitment to ribosomes upon imbibition enables rapid protein synthesis without the delay of transcription (Layat *et al*., 2014; Basbouss-Serhal *et al*., 2015; Bai *et al*., 2020). Their rapid translation during the early hours of germination is essential for germination completion and is independent of new transcription. In Arabidopsis, it has been proposed that most of the stored mRNAs in dry seeds are associated with monosomes and not polysomes, consistent with a quiescent translationally inactive state that is reactivated during germination (Bai *et al*., 2020). However, this conclusion should be interpreted with caution, as it relies on polysome profiling, a low-resolution method that cannot accurately determine ribosome occupancy or distinguish truly translationally inactive mRNAs. During subsequent seed imbibition, Basbouss-Serhal *et al*. (2015) have proposed that the 5′ untranslated regions (5′ UTRs) could regulate the translation efficiency through their GC content and the presence of upstream open reading frames (uORFs). While others proteomic and biochemical studies have also hinted at such post-transcriptional regulation (Rajjou *et al*., 2004; Galland *et al*., 2014), genome-wide analyses of ribosome occupancy in seeds are lacking, particularly for the metabolically quiescent dry state. Consequently, our understanding of how translational regulation contributes to dormancy maintenance and release remains incomplete.

Ribosome profiling (Ribo-seq) provides nucleotide-resolution maps of ribosome positions on mRNAs, allowing the identification of actively translated transcripts and features such as uORFs that modulate translational efficiency (von Arnim *et al*., 2014). uORFs are short coding sequences (often less than 50 codons in plants) located in the 5′ UTRs of mRNAs that can be translated before the main coding sequence (mORF). They modulate translation by influencing ribosome scanning and reinitiation, often repressing or fine-tuning translation of the main ORF in response to cellular or environmental cues (von Arnim *et al*., 2014). Ribo-seq in plants has been successfully used to map the translational landscape with high resolution, revealing thousands of unannotated translated ORFs (including upstream ORFs, downstream ORFs and small ORFs in non-coding RNAs), and demonstrating that many of these exert translational repression or regulatory roles (Juntawong *et al*., 2013; Hsu *et al*., 2016; Zhu *et al*., 2023). For example, in *Arabidopsis thaliana*, “super-resolution” Ribo-seq identified 7,751 unconventional translation events including ∼6,996 uORFs and 546 ORFs in presumed noncoding RNAs, some of which produce stable proteins (Wu *et al*., 2024). In rice, Ribo-seq combined with RNA-seq revealed tissue- and allele-specific translational efficiencies, detected ∼3,392 uORFs, and showed that many long non-coding RNAs are actually translated (Zhu *et al*., 2023). This approach has not yet been applied to dry seeds, a stage often presumed inactive, nor to a temporal analysis spanning dormancy release and early germination. Such analyses could reveal whether ribosomes are positioned in a “poised” state on specific transcripts, and how translational checkpoints such as uORFs shape the decision to germinate.

Here, we present the first comprehensive ribosome profiling analysis of *Arabidopsis thaliana* seeds, comparing dry dormant, stratified (non-dormant), and imbibed states. We show that dry seeds maintain an unexpectedly organized translational apparatus, with ribosomes pre-positioned at the start codons of germination-related mRNAs. We further reveal that translational efficiency patterns, and the modulation of uORF activity, distinguish dormant from non-dormant seeds and act as molecular checkpoints in the transition to germination. Our findings uncover translational regulation as a central layer of control in seed biology, complementing transcriptional programs and providing new insight into how plants fine-tune developmental transitions in response to environmental signals.

## Material and Methods

### Plant material and germination assays

*Arabidopsis thaliana*, ecotype Columbia (Col-0), seeds were obtained from plants grown at 22°C under long days (16 h of light/8 h of dark) in growth chambers as previously described (Leymarie *et al*., 2012). At harvest seeds were dormant. Dormancy was alleviated by placing seeds on a filter paper on the top of a cotton wool moistened with deionized water at 4 °C for 4 days in darkness (stratification) (Yan *et al*., 2020). Seeds, either dormant (DS) or stratified (NDS), were germinated on a layer of cotton wool covered by a filter paper sheet soaked with water for 10 d at 15 or 25°C in darkness. A seed was considered germinated when the radicle had protruded through the testa. The results presented correspond to the mean of the germination percentages obtained for 3 biological replicates of 50 seeds.

### Ribosome Profiling (Ribo-seq)

For isolation of Ribosome Protected Fragments (RPFs) ∼10 mL of pulverized frozen tissue and 30 mL of polysome extraction buffer [PEB; 200 mM Tris·HCl (pH 9.0), 200 mM KCl, 36 mM MgCl_2_, 25 mM EGTA, 5 mM DTT, 1 mM phenylmethanesulfonylfluoride, 50 μg/mL cycloheximide, 50 μg/mL chloramphenicol, 1% (v/v) Triton X-100, 1% (v/v) polyoxyethylene lauryl ether, 1% (v/v) Tween-40, 1% (v/v) Nonidet P-40, 1% (v/v) polyoxyethylene 10 tridecylether] were used to obtain the S16 fraction. RNA concentration in the S16 fraction was estimated with QUBIT HSRNA quantitation kit using a 10X diluted aliquot. RNA was digested with 50 units of RNase I per 40 µg of RNA for each sample during 1 hour at 25°C. 200µg of RNA were used for each sample. To isolate monosomes resulting from the RNAse digestion of the total population of ribosomes (i.e., monosomes and polysomes), the digested S16 fraction was layered on top of a 1.0 M sucrose cushion [400 mM Tris·HCl (pH 9.0), 200 mM KCl, 30 mM MgCl 2, 1.75 M sucrose, 5 mM DTT, 50 μg/mL chloramphenicol, 50 μg/mL cycloheximide] and centrifuged at 100000 × g for 4 h at 4 °C in a 70Ti rotor (Beckman) to obtain a polysome pellet (P100). RNA was extracted from the P100 pellet with Trizol and library preparation was performed as in Juntawong *et al*. 2013. Matching total RNA was extracted from 500µl of undigested S16 fraction using Trizol LS with the following modification. Two chloroform extractions were performed and RNA was further cleaned-up with Zymo RNA clean and concentrator columns. RNA seq libraries were constructed with ScriptSeq™ v2 RNA-Seq (Epicentre) following the manufacturer instructions.

### Short Read Processing, Quality Assessment, and Alignment

FASTQ files were obtained with the base caller provided by the instrument. All data analysis steps were performed using a combination of command-line software tools and the R packages from Bioconductor including the SystemPipeR workflow for ribo-seq. The Short-Read package was used for adaptor trimming, quality assessment, and filtering of reads (90). Adaptors were removed with the FASTX-Toolkit program Fastx-Clipper using the following adaptor sequences: ribo-seq, AGATCGGAAGAGCACACGTCTGAACTCCAGTCA; mRNA-seq, CTGTAGGCACCATCAATAGATCGGAAGAG. Comprehensive quality reports of raw and trimmed read sets were generated with the seeFastq from SystemPipeR package. For polyA+-RNA-seq, ribo-seq the trimmed and quality filtered reads were mapped with the splice junction aware short read alignment suite Tophat, version 2.0.1, to the Arabidopsis genome sequence of The Arabidopsis Information Resource, release version 10, allowing only unique alignments (i.e., each read was allowed to map to one location only) with ≤2-nt mismatches.

Expression analyses were performed by generating read count data for various feature types of interest (e.g., exons-by-genes, CDSs, UTRs, exons) using the summarizeOverlaps function from the GenomicRanges package. The minimum fold change and false-discovery rates (FDRs) used by this study as confidence filters are given in Results. uORFs were predicted for 5′-UTR sequences from gene annotation of Araport11 GFF file (86) using the predORF function from systemPipeR (91) with ATG as start codon and a minimum length of 60 nt. Expression differences among uORF and mORF features were determined as in (Ge *et al*., 2020)Juntawong *et al.,* 2013). Only uORFs with the highest uORF vs. mORF ratio were retained for the analysis. For protein-coding genes, translational efficiency (TE) was calculated comparing rpkM values of ribosome-protected fragments with mRNA rpkM values for the coding sequence (excluding UTRs) for genes with a minimum of 5 rpkM in the RF and RNA-seq datasets for all conditions.

Differential TE analysis was performed using using the XTail pipeline (Xiao *et al*., 2016). Metagene analysis of RPF positioning around Start and Stop codon and 3’nt periodicity analysis were performed using Plastid (Dunn & Weissman, 2016).

Heatmaps in were performed with SRPLOT (Tang *et al*., 2023). Hierarchical clustering and heat map of TE and gene expression was performed using the Pheatmap R package using a z-score normalization. Gene Ontology (GO) analysis was performed with ShinyGO V0.80 (Ge *et al*., 2020) and agriGO v2.0 (Tian *et al*., 2017).

### Data Availability and Analysis

The RNA-seq data generated in this study have been deposited in the [NCBI Gene Expression Omnibus (GEO)](https://www.ncbi.nlm.nih.gov/geo/). Raw sequencing reads (FASTQ files) and processed count matrices are available.

### Cloning of *MARD1* and *PAO4* uORF

To generate the *35SuORF-MARD1::LUC* and *35SuORF-PAO4::LUC* constructs, the sequences before the mORF of *MARD1* (-272bp, *AT3G63210*) and *PAO4* (-391bp, *AT1G65840*) were amplified using DreamTaq DNA Polymerase (Thermo) from genomic Col-0 DNA. The *uORF-MARD1* and *uORF-PAO4* PCR fragments were digested with *SpeI-NotI* and *SpeI-NcoI*, respectively; and ligated into a modified *pGreen800-35S::LUC* vector (kindly provided by Juliette Puyabert). A site-directed mutagenesis strategy was used to modify the ATG codon of the uORFs on *35SuORF-MARD1::LUC* and *35SuORF-PAO4::LUC* constructs. To create the specific mutations, a PCR with the Phusion High-Fidelity DNA Polymerase (Thermo) was performed with specific primers followed by a removal of the template DNA (DpnI enzyme, Thermo), a 5’ phosphorylation (T4 Polynucleotide Kinase, Thermo) and the ligation (T4 DNA Ligase, Thermo) before transformation of *E. coli* to generate *35Smut-uORF-MARD1::LUC* and *35Smut-uORF-PAO4::LUC* constructs. The primers used for amplification of the uORF and the mut-uORF are described Table S1. All plasmids were checked by sequencing.

### Protoplast Isolation, Transfection and Dual Luciferase Assays

Arabidopsis mesophyll protoplasts were isolated as described by Yoo *et al.,* 2007. In brief, leaves of 4-week-old Col-0 plants grown in short photoperiod (8h light/16h dark) were collected and cut in 0.5–1-mm leaf strips. Leaf strips were dipped into an enzyme solution (20 mM MES pH 5.7, 1.5% wt/vol cellulase R10, 0.4% wt/vol macerozyme R10, 0.4 M mannitol, 20 mM KCl, 10 mM CaCl_2_, 1–5 mM b-mercaptoethanol and 0.1% BSA). After vacuum for 30 min, digestion was maintained for 3h in dark at RT. Reaction was stopped by adding an equal volume of W5 solution (2 mM MES pH 5.7, 154 mM NaCl, 125 mM CaCl2 and 5 mM KCl) and filtered through a 100-mm nylon mesh. Released protoplasts were resuspended in the MMG solution (4 mM MES pH 5.7, 0.4 M mannitol and 15 mM MgCl2). Around 2x10^4^ protoplasts were mixed with 10 µg of plasmid DNA per construct, and mixed with the PEG solution (30% PEG 4000, 0.2 M mannitol and 100 mM CaCl2) and incubated for 10 min at RT. Protoplasts were then resuspended in the WI solution (4 mM MES pH 5.7, 0.5 M mannitol and 20 mM KCl) and incubated for 16 h in dark at RT. Protoplasts were then harvested by spinning at 1,000 x g for 30 s.

The Dual-Luciferase Reporter Assay System (Promega) was performed according to manufactures instructions. In brief, protoplasts were resuspended in 1x passive lysis buffer (Promega) and briefly vortexed. Lysed protoplasts were mixed then with the Dual-Glo® Luciferase Reagent. After incubation for 15min at RT, firefly luciferase activity (fLUC) was measured with a Microplate reader (TECAN). Then, the Renilla luciferase activity (rLUC) was measured by adding the Dual-Glo® Stop & Glo® Reagent. Luminescence was measured in technical triplicates for all combinations of transfected plasmids.

## Results

### Seed germination and dynamics of transcriptome and ribosomal association

At harvest Arabidopsis seeds were dormant, i.e. they germinated poorly at 25°C in the darkness (Fig. 1A) but fully germinated at 15°C (data not shown). Four days of stratification fully released seed dormancy and stratified seeds became able to fully germinate after 6 days in these conditions (Fig. 1A). Based on this biological model, we explored the involvement of transitional efficiency in the regulation of seed germination in dormant (dry seeds) and non-dormant (stratified) seeds and at different time points of their imbibition at 25°C, before radicle protrusion (Fig. 1B). We performed both RNA-seq and Ribo-seq on dormant (DS) and non-dormant (NDS) seeds at different hours after imbibition at 25°C (0, 3, 15 and 24 hai). Distribution of the RNA-seq and Ribo-seq samples is shown in the Principal Component Analysis (PCA), in which PC1 and PC2 explained 64% of the total variance (Fig. 1C). The analysis clearly separated dormant from non-dormant stratified seeds, either for transcripts abundance (RNA-seq) or ribosome-protected fragments (RPFs) (Ribo-seq) (Fig. 1C). It also shows the dynamics of the transcriptome and of the translational landscape during seed imbibition since both strongly differed from 3 to 24 hai in dormant seeds and in a much lesser extent in stratified seeds (Fig. 1C, Dataset S1). The PCA Fig. S1A also shows a clear separation of samples by time point and sequencing method, indicating strong temporal dynamics and distinct profiles between RNA-seq and Ribo-seq datasets. The correlation tree and replicate reproducibility plots further confirm high consistency across samples, supporting the robustness of the dataset for downstream analyses (Figs S1A and S1B). In addition, Ribo-seq libraries showed a strong and consistent enrichment of reads on annotated CDS across all samples, with minimal contribution from non-coding features, confirming high translational specificity of the datasets. Sense reads largely dominate over antisense signals, indicative of low background and limited off-target priming. The residual mapping to UTRs and other genomic compartments remains within expected bounds for high-quality plant Ribo-seq libraries (Fig. S2).

**Figure 1.**
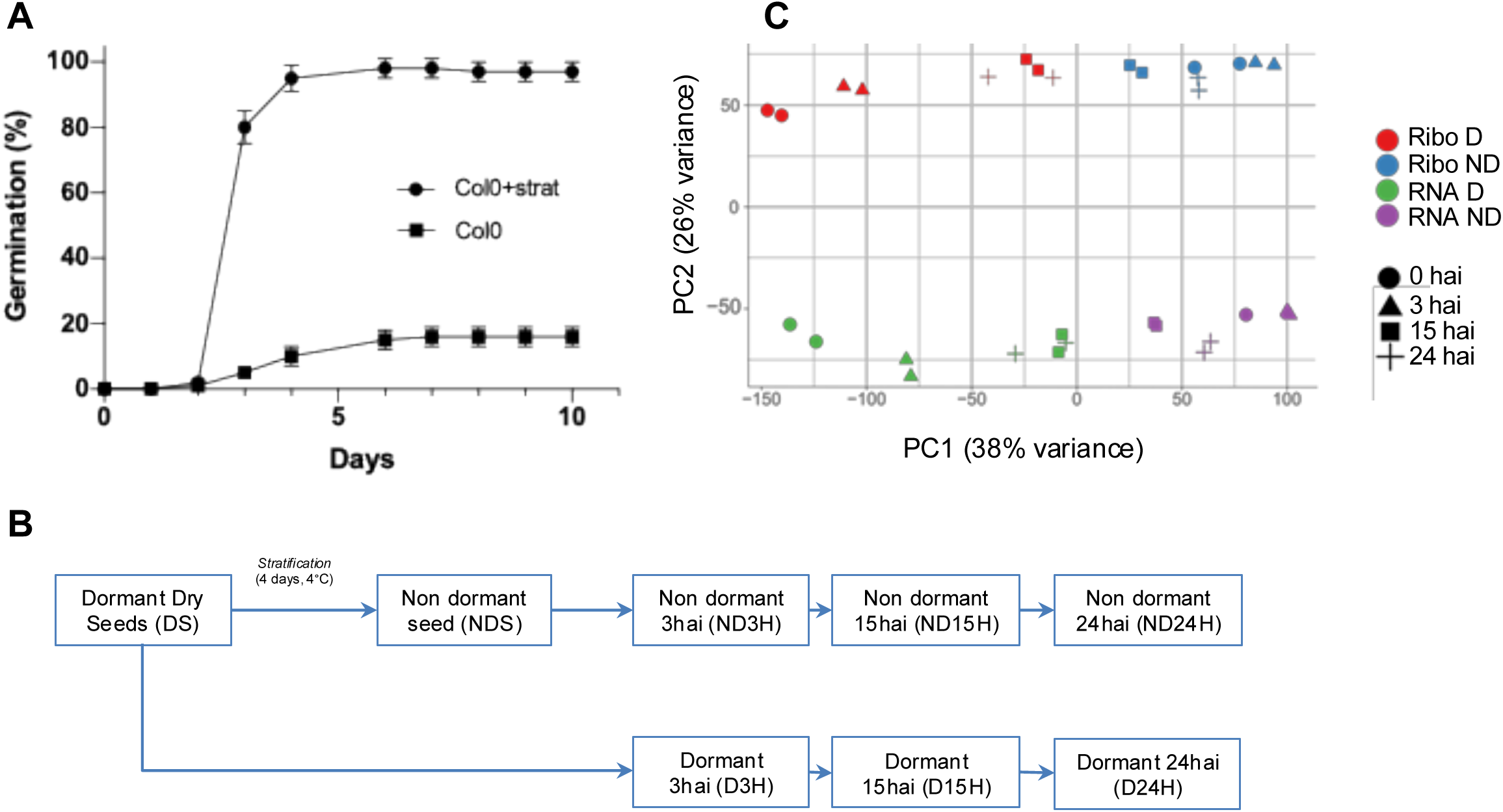
Dormant and non-dormant stratified seeds have differential germination phenotypes. **A)** Germination assay of freshly harvested dormant Col-0 seed and 4 days cold treated Col-0 seeds (Col-0 + Strat) at 25°C in darkness. Means of 3 replicates ± SD. **B)** Experimental design to produce Ribo-seq and RNA-seq samples. **C)** PCA analysis of the Ribo seq (Ribo) and RNA-seq (RNA) seed samples (D, dormant; ND, non-dormant) at 0, 3, 15 and 24 hours after imbibition (hai).

We first compared transcript abundance and translation in dry and stratified seeds (0H_DS vs NDS), thus underlying the processes associated with dormancy release (Fig. 2A). Among the 13508 genes detected in both datasets, we identified 1873 that were regulated similarly at the transcriptional and translational levels (homodirectional) (Dataset S1). This list includes genes on which mRNA level and ribosome footprint density increase and genes on which mRNA and ribosome footprints decrease in DS vs NDS; suggesting that transcript availability of those genes directly drives protein synthesis. We also identified 631 genes that were regulated only at the transcriptional level (transcription only) and 1298 only at the translational level (translation only) (Fig. 2A; Datasets S1 and S2). A reduced number of genes (only 66) were regulated in an opposite way (opposite change), indicating post-transcriptional/translational rather than transcriptional control (Fig. 2A; Datasets S1 and S2). A relatively similar relationship between transcription and translation in dormant and non-dormant stratified seeds was evidenced at 3 hai (3H_DvsND; Fig 2A) where the large majority of differentially expressed genes (831) shown homodirectional changes and only 182 genes were only translationally regulated (Datasets S1 and S2). In contrast, further imbibition (15 and 24 hai) drastically reduced changes in transcriptional and translation regulation between dormant and non-dormant seeds (Fig 2A, Datasets S1 and S2). Gene Ontology (GO) term enrichment analysis of genes homodirectionally regulated was evaluated, independently of seed status or duration of imbibition (Dataset S3). Genes involved in response to stress, hormones and environmental signals appeared as being repressed (down) at the level of the transcript abundance and translation (Fig. 2B). In contrast, mRNAs involved in translation and protein synthesis were up regulated at both levels (Fig. 2C). It is worth noting that another set of genes specifically associated with translation and RNA processing were also regulated at the level of translation only (Dataset S3). The genes that were up or down regulated only at the transcriptional level cover various GO categories that do not seem to be closely to the germination and dormancy processes (Dataset S3)

**Figure 2.**
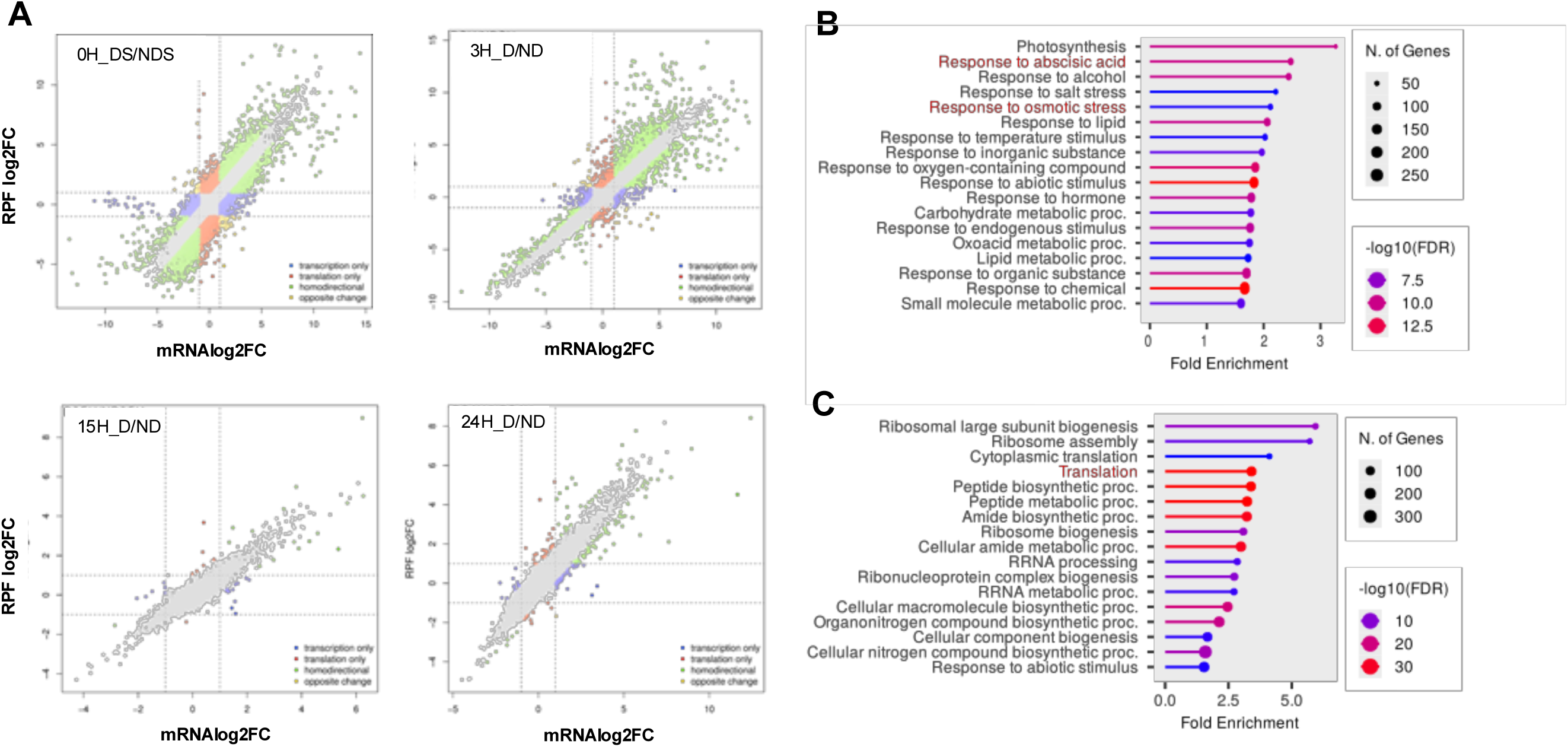
Dynamics of transcriptome and ribosomal association. **A)** Log2 fold change (log2FC) of RPF abundance and RNA-seq reads abundance for genes between dormant and non-dormant seeds (DS, NDS); dormant (D) and non-dormant (ND) seeds after 3 h, 15 and 24 hours after imbibition. The color of the dot classifies genes according to their mode of regulation. **B)** GO analysis of genes homodirectionally downregulated and **C**) upregulated during stratification (all sample times combined, 0 h, 3 h, 15 h and 24 h).

We investigated the dynamics of ribosome occupancy of a set of well-described regulators of seed germination during stratification and imbibition of dormant and non-dormant seeds (Fig. 3A, Dataset S4). Two major clusters separate dormancy from germination gene markers. The first major cluster is divided in a subcluster including transcripts being positively associated with dormancy, such as *ABA1* (Lopez-Molina *et al*., 2002), *ABI5* (Lopez-Molina *et al*., 2001) or *TT4* (Debeaujon & Koornneef, 2000). This subset of genes was highly associated with ribosomes in dry dormant seeds and seed stratification markedly decreased ribosome occupancy for these transcripts. The other subcluster contained other transcripts being considered as major regulators of seed dormancy, including for example *DOG1* (Graeber *et al*., 2013) or *NCED6* (Lefebvre *et al*., 2006). They were also covered by ribosomes in dry dormant seeds but the dormancy release treatment per se (stratification) did not modify markedly this feature, however their coverage by ribosomes subsequently decreased dramatically during seed imbibition (Fig. 3A). The heatmap also shows that maintenance of dormancy mostly relied on the translation of negative regulators of germination during early imbibition (3 hai). For example, the ribosome occupancy of *DOG1* was high in dry dormant seeds and 3 h of imbibition, but strongly decrease after 15 h of imbibition (Fig. 3A). On the second major cluster, in contrast, the ribosome occupancy of transcripts associated with germination, ie. *GA3ox2* or *PhyA*, was low in dormant seeds but was significantly high after seed dormancy release (stratified), remaining constantly high untill 24 h of imbibition (Fig. 3A). The coverage maps of RPFs clearly indicate their activate translation as a function of seed physiological status, evidencing by a strong ribosome occupancy on dormant (*DOG1* and *ABI5*, Fig. 3B) and germination (*PHYA* and *GA3ox2*, Fig. 3C) gene markers.

**Figure 3.**
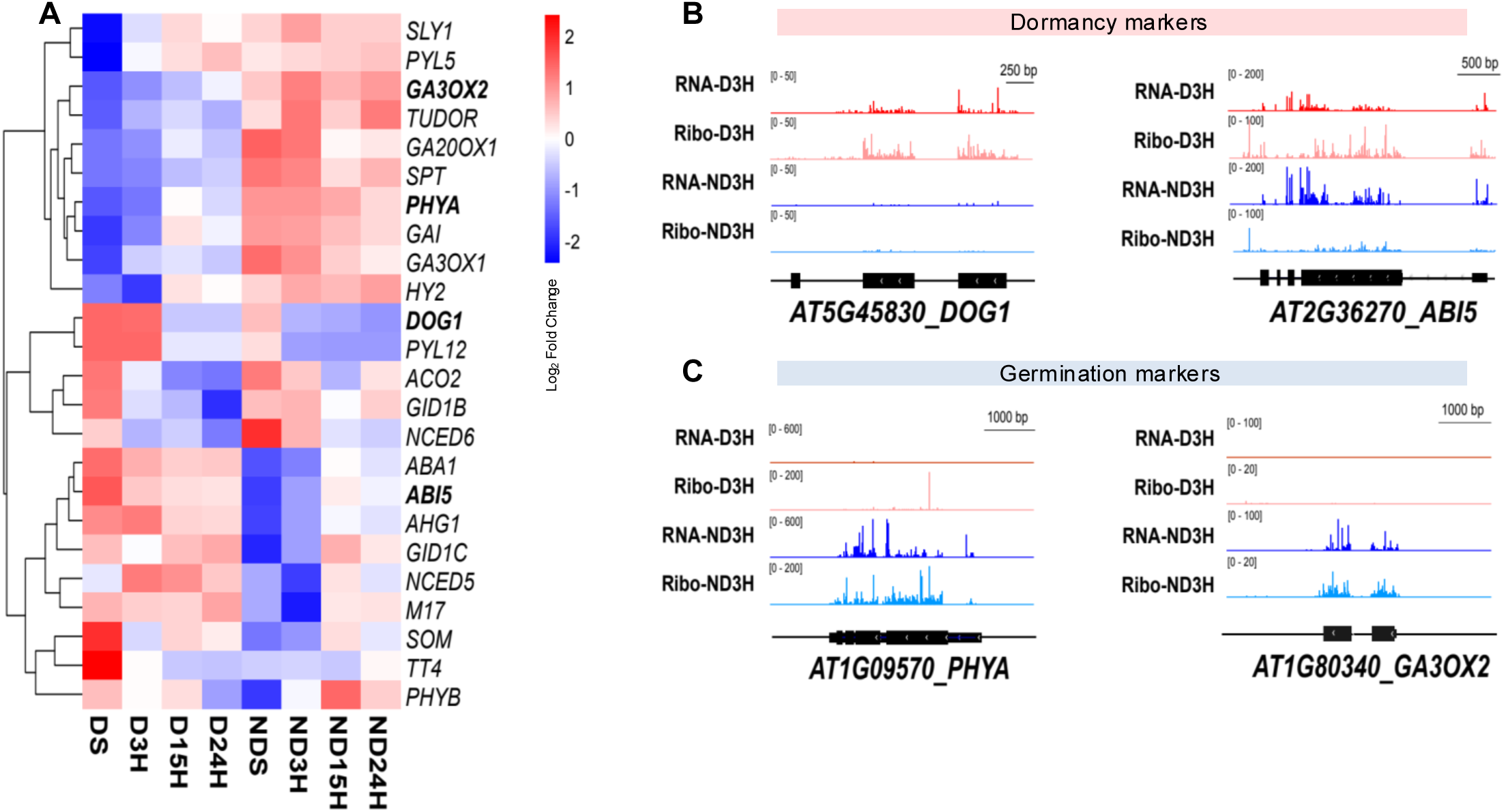
Dynamics of ribosome occupancy during seed dormancy and germination. **A)** Heatmap and hierarchical clustering of log2 Ribo-seq reads abundance between dormant and non-dormant seeds (DS, NDS); dormant and non-dormant seeds after 3 h (D3H, ND3H), 15 (D15H, ND15H) and 24 (D24H, ND24H) hours after imbibition. Coverage plots for ribosome-protected fragments (RPFs) and RNA-seq reads data mapped for dormancy **(B)** and germination **(C)** gene markers. The gene models for *DOG1* (*DELAY OF GERMINATION 1*), *ABI5* (*ABA INSENSITIVE 5)*, *PHYA (Phytochrome A), GA3OX2 (GIBBERELLIN-3-OXIDASE 2)* genes are displayed in black below the coverage plots.

### Changes in translational efficiency

Ribo-seq allowed the evaluation of translational efficiency (TE) for individual mRNAs, by calculating the ratio of RPF reads relative to the density of mRNA fragments in the CDS (Brar & Weissman, 2015). In order to investigate the relevance of ribosome loading of seed stored mRNA on seed dormancy and germination, we grouped transcripts in deciles according to their ribosome density in dry seeds and investigated their change during imbibition in dormant and non-dormant seeds (Fig. 4A and B, Fig. S3, Dataset S5).

**Figure 4.**
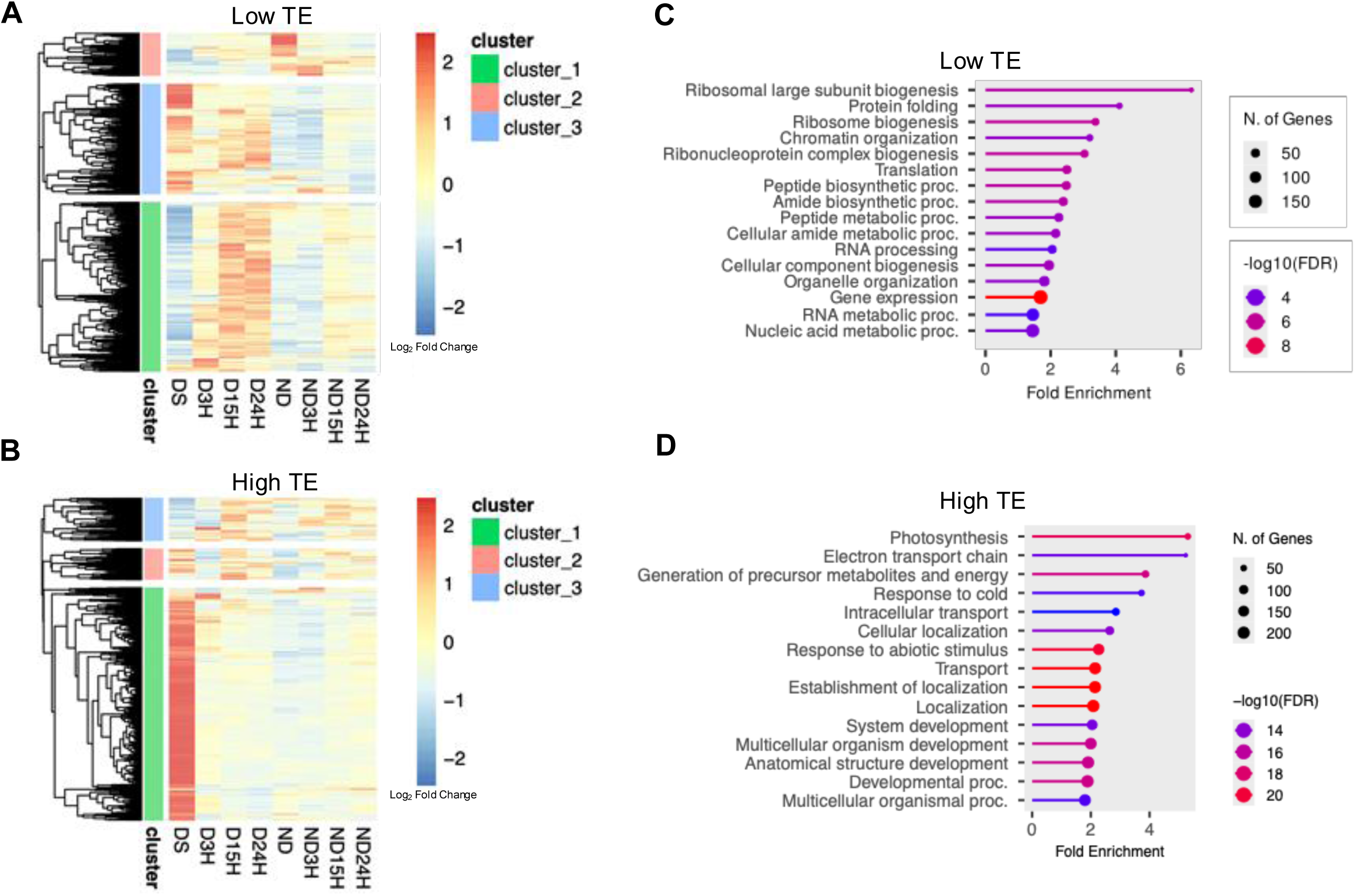
Average translation efficiency (TE) of genes in each sample. **A**) Hierarchical clustering of TE during imbibition of dormant and non-dormant seeds (DS, NDS); dormant and non-dormant seeds after 3 h (D3H, ND3H), 15 (D15H, ND15H) and 24 (D24H, ND24H) hours after imbibition for genes with low TE (1st decile) or **B**) high TE (10th decile) in dry seeds (DS). **C**) GO analysis of cluster 1 from low TE and **D)** cluster 1 from high TE. x axis represent the *p-*value (-log10) of enrichment (hypergeometric test) for each GO categories. Only non-redundant significant categories are represented.

The distribution of all detected mRNAs with a low TE (1^st^ decile) in dry seeds is shown in Fig. 4A. Interestingly, we found that TE was globally lower in non-dormant seeds than in dormant seeds, at each time point of imbibition (Fig. 4A), suggesting that translational activity is more involved in dormancy maintenance than in germination completion. Moreover, average TE was significantly higher in dry seeds than in imbibed dormant and non-dormant seeds, suggesting an important loading of seed stored mRNA into ribosomal complexes during seed maturation. Cluster 1 from the low TE group, displayed transcripts with an increasing TE in dormant imbibed samples (Fig. 4A, Dataset S5). It was significantly enriched in GO terms related to translation, ribosome biogenesis, and protein folding, reflecting a preparatory role for protein synthesis machinery that may be primed for rapid activation in dormant seeds (Fig. 4C).

We then investigated TE evolution of transcripts that displayed a high TE in dry seeds (10^th^ decile, Fig. 4B, Dataset S5). Genes with high initial TE (Fig. 4B) showed more variable behavior, with some gaining TE over time (Cluster 3), especially in dormant seeds, possibly reflecting delayed or tightly regulated translation initiation under dormancy conditions. The TE of a large majority of these mRNAs further decrease during stratification and imbibition, whatever seed were dormant or not (Cluster 1, Fig. 4B). The GO analysis of this cluster covered a wide range of developmental/cellular processes without a strong common identity (Fig. 4D). The changes in TE of transcripts belonging to other clusters were less consistent but the general trend was that mRNAs displaying high TE in dry seeds were then sufficient to initiate protein production during seed imbibition.

### Translation dynamics during seed imbibition

Because the ribosome does not protect the entire mRNA equally, the exact position of the ribosome on the transcript is inferred by identifying the P-site, which corresponds to the codon being actively translated. For footprints of 28–30 nucleotides, the P-site is located at a fixed distance from the 5′ end of the read (here, an offset of 14 nucleotides). Mapping these inferred P-site positions along the 5′ UTRs, coding sequences (ORFs), and 3′ UTRs therefore reveals where ribosomes are positioned across transcripts and allows assessment of translation initiation, elongation, and non-canonical ribosome occupancy. Here, to gain further insight into ribosomes dynamics during seed germination, we analyzed ribosome positioning across mRNAs by mapping the relative locations of ribosome P-sites from ribosome footprint reads (offset of 14 nt, footprints 28-30nt long) along the 5′ UTRs, ORFs, and 3′ UTRs of all detected transcripts with a minimum of 5 TPM RPF reads in all samples (Fig. 5). In dry dormant seeds, we observed a characteristic peak centered at the start codon, indicating that a subset of stored seed mRNAs is already engaged in translation initiation. Within the CDS, ribosomes showed significant enrichment of P-sites in the coding frame and exhibited a clear 3-nt periodicity, suggesting that desiccated seeds retain elongating ribosomes on coding regions. Consistent with observations in other plant tissues, RPF density across the 3′ UTRs was very low in all samples. As expected, dry seeds lacked the typical peak just upstream of the stop codon-a hallmark of active translation elongation. However, a distinct peak at the stop codon became apparent 3 hours after imbibition (D3H; Fig. 5A), confirming previous findings that active translation elongation resumes early during imbibition. Inset in Fig. 5A summarizes the translational state we demonstrate in dry seeds, in which ribosome profiling reveals strong P-site enrichment at annotated start codons. In this model initiating ribosomes are stalled at the AUG, with the initiator Met-tRNAᵢ positioned in the P-site while the A-site remains unoccupied, awaiting binding of the aminoacyl-tRNA corresponding to the second codon.

**Figure 5.**
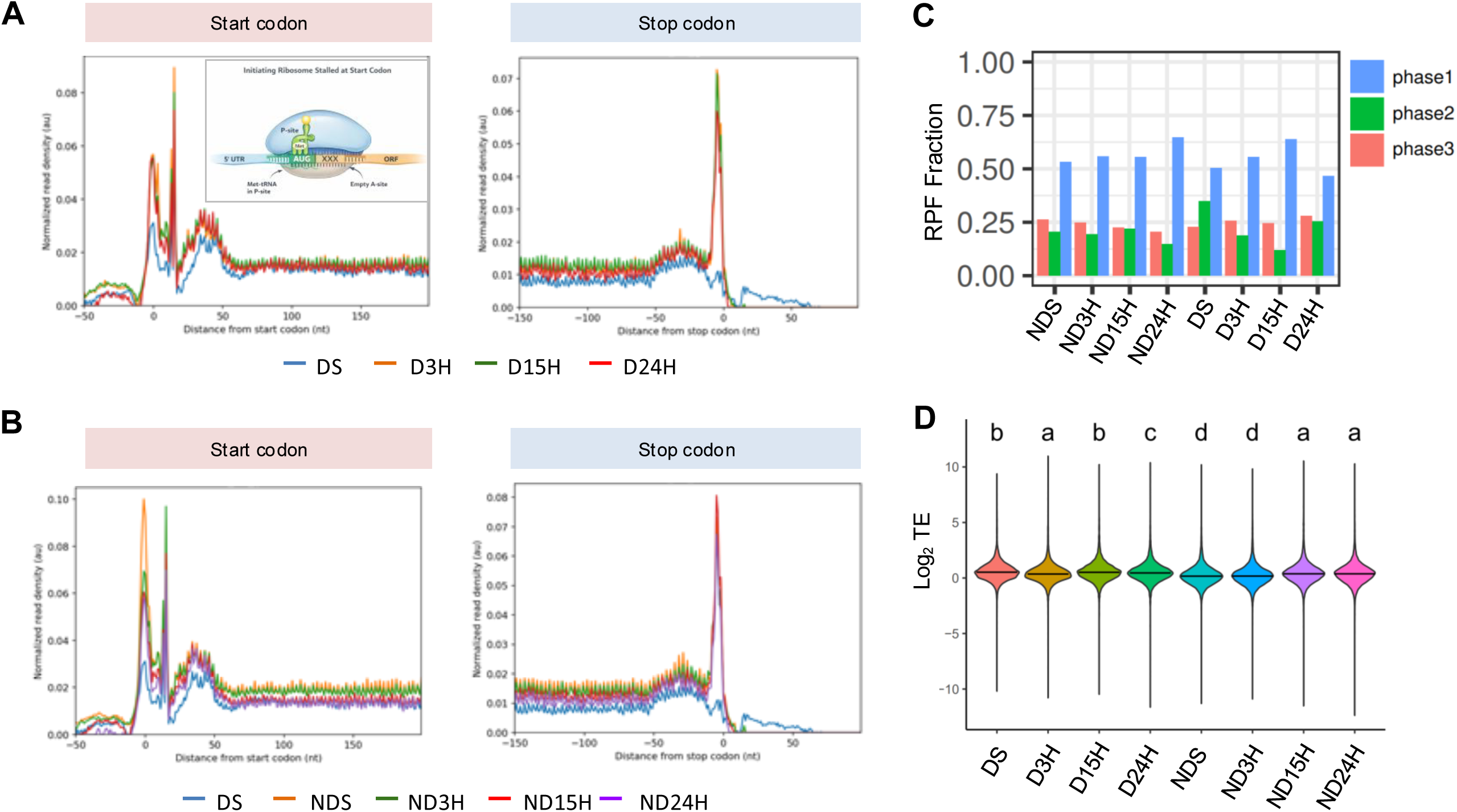
Dynamics of ribosome-protected fragments (RPF) distribution during seed germination and dormancy. Metagene analysis of P-Sites distribution around start and stop codon on all detected gene during imbibition of dormant (**A)** and non-dormant seeds (**B)**. **C)** P-Sites enrichment analysis on the 3 coding phase of each CDS (phase1 correspond to the right coding phase)**. D**) Distribution of log2 transformed TE value for each time points. Significant differences are indicated with letters (Statistical differences between groups were assessed using one-way ANOVA followed by Tukey’s post-hoc test for multiple comparisons. Groups sharing the same letter are not significantly different, whereas groups with different letters are significantly different p.adjusted < 0.05). Dormant and non-dormant seeds (DS, NDS); dormant and non-dormant seeds after 3 h (D3H, ND3H), 15 (D15H, ND15H) and 24 (D24H, ND24H) hours after imbibition. Inset Fig. 5A shows a schematic representation of an initiating ribosome stalled at the start codon in dry seeds.The initiator Met-tRNAᵢ is bound in the P-site, while the E-site is empty and the A-site has not yet accommodated the incoming aminoacyl-tRNA corresponding to the second codon.

Notably, while both dormant (Fig. 5A) and non-dormant (Fig. 5B) seeds showed conserved global translation patterns, the magnitude and sharpness of the peaks differed over time, suggesting global modulation of translation (TE) during imbibition. After 3h of seed imbibition of dormant seeds at 25°C, ribosome occupancy increased around start and stop codon but did not change in later time points. In non-dormant seeds, RPF density peaks specifically at the start codon after stratification (ND) suggesting a positive effect of stratification on global translation and particularly on the dynamics of translational initiation. However, this signal gradually decreased during imbibition of non-dormant seeds (ND3, ND15, ND24), indicating downregulation of global translation initiation as seeds progressed through early germination.

P-site enrichment across the three coding phases reflects the 3-nt periodicity of translating ribosomes, with phase 1 corresponding to the annotated coding frame, while phases 2 and 3 represent the second and third nucleotide positions of codons and serve as indicators of data quality and translational frame fidelity. Here, P-site enrichment analysis revealed a strong triplet periodicity across the 3 coding frames for all conditions, with a clear enrichment in phase 1 corresponding to the correct coding frame, confirming high-quality ribosome profiling data and accurate assignment of ribosome footprints to actively translating ribosomes (Fig. 5C). This periodicity was conserved across dormant and non-dormant seeds and throughout imbibition, indicating preserved elongation fidelity during the transition from dormancy to germination. Analysis of translational efficiency (TE) distributions showed significant, time-dependent differences between physiological states, with globally higher TE in dormant seeds that progressively decreased during imbibition, as supported by one-way ANOVA followed by Tukey’s post-hoc test (p.adjusted < 0.05) (Fig. 5D). These global changes in TE prompted us to examine whether gene-specific regulatory mechanisms, particularly upstream open reading frames (uORFs), contribute to the selective modulation of translation during dormancy release and early germination.

### uORF mediated regulation contribute to translation selectivity during seed imbibition and dormancy

Putative translated uORF were extracted from (Wu *et al*., 2024) and further filtered for the presence of RPF mapping on their respective location in our datasets. To further understand the translational control exerted by upstream open reading frames (uORFs), we quantified the ratio of translational efficiency between main ORFs (mORFs) and uORFs across all samples (Fig. 6A). The distribution of log₂-transformed mORF/uORF ratios of all transcripts revealed marked variation, with a general trend toward decreased mORF translation relative to uORFs in dormant and non-dormant seeds, particularly at 15 h of imbibition. This shift suggests translational repression via uORFs, potentially inhibiting the expression of proteins required for germination or dormancy maintenance. Hierarchical clustering based on mORF-to-uORF ratios (Fig. 6B, Dataset S6) further highlighted distinct translational landscapes between dormant and non-dormant seeds. It is worth noting that stratification (DS vs NDS) resulted in either increase or decrease in on mORF-to-uORF ratios in a similar proportion (Fig. 6B). Dormant seed samples (DS, D3H, D15H, D24H) formed a separate cluster from non-dormant samples (NDS, ND3H, ND15H, ND24H), indicating consistent differences in uORF-mediated translational control. The separation was most pronounced at early time points, suggesting that differential translation via uORFs may play a role in establishing or maintaining dormancy status during the early phase of water uptake. Together, these data point to a dynamic reprogramming of translation during seed imbibition, in which uORF regulation contributes to the differential expression of proteins in dormant versus non-dormant states.

**Figure 6.**
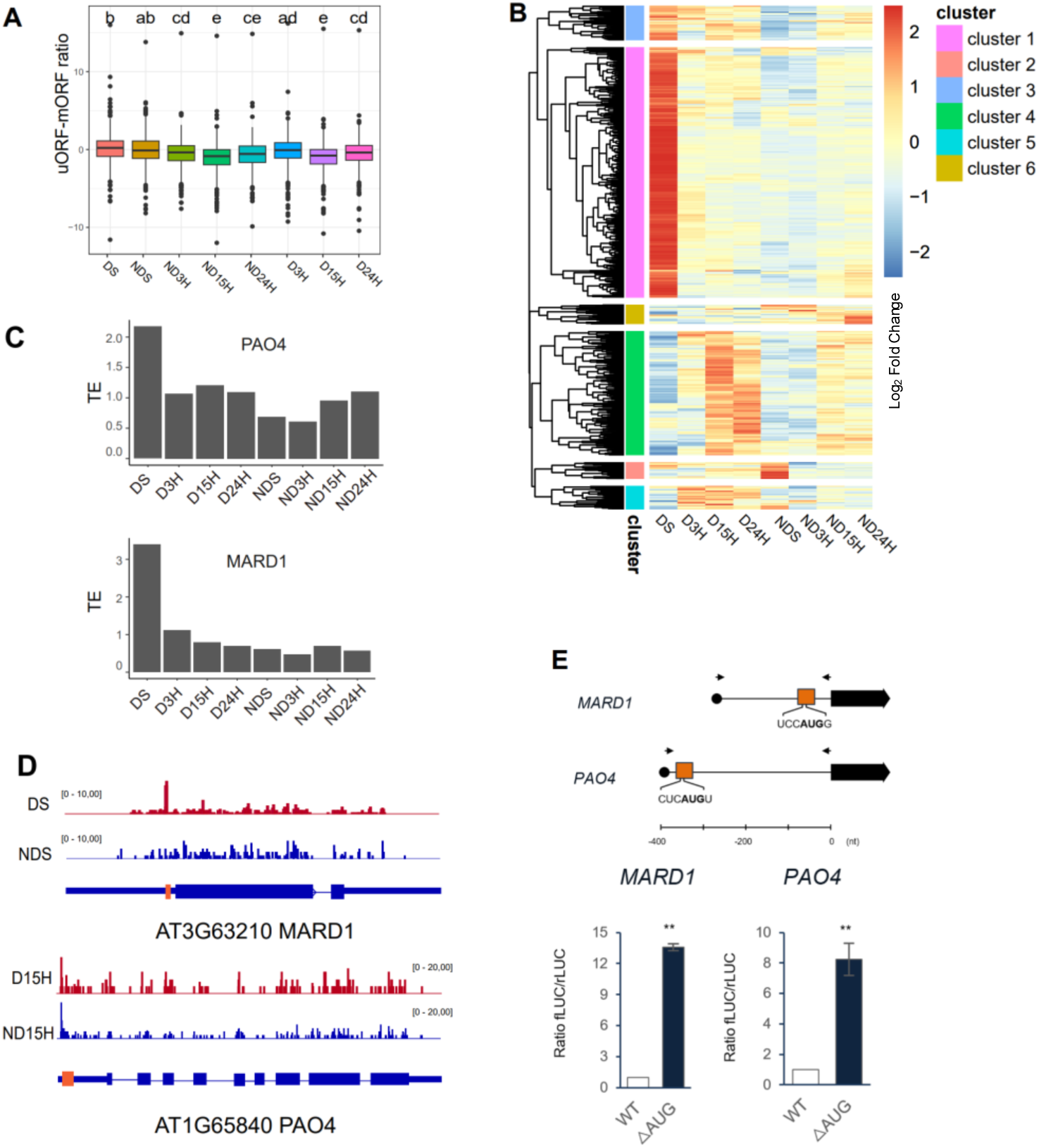
uORF mediated regulation during seed dormancy and germination. **A**) Global distribution and **B)** hierarchical clustering of uORF/mORF ratio for all translated uORF in DTE genes. **C**) TE and uORF/mORF ratio evolution of *PAO4 and MARD1* genes **D**) Coverage plots for ribosome-protected fragments (RPFs) data mapped to *MARD1* and *PAO4* genes. Genes are displayed in black below the coverage plots. Orange box indicates predicted uORF. **C**) Schematic representation of the 5′-UTRs containing conserved AUG uORFs of *MARD1* and PAO4 genes used for transient expression assay. **E**) Each reporter plasmid containing the WT or ΔAUG uORF version of *MARD1* and *PAO4* genes were transfected into Arabidopsis protoplasts. Firefly luc (fLUC) activity was normalized to Renilla luc (rLUC) activity. Data represent the mean ± SD of the normalized LUC activities (fLUC/rLUC) of two biological replicates, relative to that of the corresponding WT reporter construct. Double asterisks indicate significant differences between the WT and ΔAUG -mutant constructs at *p* < 0.01, respectively, as determined by Student’ s t-test.

To investigate the functional role of uORFs in translational regulation during seed imbibition, we focused on two candidate genes known to be involved in dormancy and stress response pathways: *MEDIATOR OF ABA-REGULATED DORMANCY 1* (*MARD1*) and *POLYAMINE OXIDASE 4* (*PAO4*). *MARD1* codes for a zinc-finger transcription factor, and is an important downstream component of the ABA signaling pathway that mediates ABA-regulated seed dormancy in Arabidopsis. The *mard1* seeds are less dormant and germinate in total darkness (He & Gan, 2004). PAO4 is an enzyme responsible for polyamine catabolism. Polyamines are nitrogen-containing compounds crucial to maintain control of the cellular homeostasis of Reactive Oxygen Species (Benkő *et al*., 2022).

Temporal profiles of translational efficiency (TE) and mORF-to-uORF ratio for these two transcripts across timepoints of dormant and non-dormant seeds are depicted in Fig. 6C. In dry dormant seeds, *MARD1* displayed a high TE compared to non-dormant seeds, which dramatically decrease in imbibed seeds. In contrast, *PAO4* showed a modest but progressive increase in TE after imbibition, particularly in non-dormant seeds. Both genes harbor conserved AUG-initiated uORFs in their 5′ untranslated regions (5′-UTRs) (highlighted in orange boxes in Fig. 6E). Notably, RPF coverage plots confirmed active ribosome occupancy over the uORFs for both genes, suggesting that these uORFs are translated and likely contribute to translational repression of the downstream coding sequence at different time points (Fig 6D). To validate the regulatory role of these uORFs experimentally, we conducted transient expression assays in *Arabidopsis* protoplasts using luciferase reporter constructs (Fig. 6E). Wild-type (WT) constructs containing the native 5′-UTRs of *MARD1* or *PAO4* were compared to ΔAUG variants in which the conserved start codons of the uORFs were mutated. The ΔAUG mutants displayed a significant increase in luciferase activity compared to their WT counterparts (*p* < 0.01, Student’s *t*-test), confirming that these uORFs exert a repressive effect on translation. This repression was stronger in *MARD1* than *PAO4* (ratio fLUC/rLUC), in agreement with the *in vivo* RPF and TE data. These results demonstrate that the uORFs in *MARD1* and *PAO4* act as translational repressors, and that their regulatory impact is dynamically modulated during seed imbibition, particularly in relation to dormancy status. This supports a broader model in which uORF-mediated translational control serves as a post-transcriptional mechanism for fine-tuning gene expression in the early stages of seed germination.

## Discussion

Seed germination marks a decisive developmental switch governed by a cascade of molecular events, many of which occur post-transcriptionally. While transcriptional programs underlying dormancy and germination have been extensively characterized (Bassel *et al*., 2011; Narsai *et al*., 2017), our study reveals that translational regulation, and in particular the positioning and activity of ribosomes on key mRNAs, is a central, previously underappreciated layer of control. Through the use of ribosome footprint (Ribo-seq) in *Arabidopsis thaliana* seeds, we provide the first comprehensive description of ribosome occupancy in dry seeds and demonstrate that ribosomes are actively pre-positioned at the start codon and coding sequences (CDS) of key transcripts, reflecting a state of translational readiness even before imbibition begins.

### Translational control is a major regulator of germination

Stratification is routinely used to overcome the physiological barriers that prevent Arabidopsis seed germination, and 4 d of this treatment fully released seed dormancy (Fig. 1A). Cold temperatures experienced during stratification are known to trigger biochemical changes within the seed that deactivate dormancy mechanisms and typically involve the breakdown of inhibitors or the activation of enzymes that promote germination processes (Yamauchi *et al*., 2004) and are associated with in depth transcriptome reprogramming (Narsai *et al*., 2011). In this former study the abundance of more than 7000 transcripts increased over 2 days of stratification which is roughly half of the number of transcripts identified in the present study after 4 days of stratification, as shown by the PCA in Fig. 1C. Similar discrepancy between dry dormant and stratified seeds was also evidenced at the level of the ribosome associated transcripts (Fig. 1C). The PCA showed that differences between samples, either at the RNA-seq or Ribo-seq levels, tended to dramatically decrease during seed imbibition suggesting that the molecular changes that govern the decision to germinate take place during stratification and early imbibition. (Kimura & Nambara, 2010) demonstrated that an active resumption of transcription and of mRNA was initiated in seeds imbibed for 30 min. The effect of stratification on seed translational activity is less documented but Arc *et al*. (2012) showed that the abundance of ca. 400 proteins changed during 5 days of this treatment which is far less than the number of transcripts identified in our Ribo-seq data.

Our comparative analysis of transcript abundance and translational activity between dry and stratified seeds reveals distinct regulatory landscapes that underlie dormancy release. The observation that 1,873 genes were regulated in a homodirectional manner at both the transcriptional and translational levels suggest a coordinated response governing early seed metabolic reactivation. This pattern is consistent with previous findings that a subset of stored mRNAs in dry seeds is translationally primed for rapid activation upon imbibition (Basbouss-Serhal *et al*., 2015; Bai *et al*., 2020). Importantly, the identification of 1,298 genes showing an exclusive regulation at the translational level highlights the significance of post-transcriptional control during dormancy release. Such regulation may reflect selective ribosome loading onto pre-existing mRNAs, a mechanism known to occur in seeds where global translation is generally repressed yet specific transcripts escape this inhibition (Galland *et al*., 2014; Layat *et al*., 2014). Ribosome-associated mRNA pools in dry seeds, enriched for transcripts related to translation, protein folding, and hormonal signaling, are thought to enable rapid proteome reprogramming during early germination (Bai *et al*., 2020). At 3 hai, a comparable transcription-translation relationship was observed between dormant and non-dormant stratified seeds, with most differentially expressed genes (DEGs) showing concordant regulation. This early stage appears to be marked by general reactivation of cellular machinery regardless of dormancy status. However, the observation that only 182 genes were translationally regulated at this time point in a dormancy-specific manner may reflect subtle early decisions that lead to divergent developmental trajectories. Interestingly, as imbibition progressed, differences between dormant and non-dormant seeds diminished in terms of both transcriptional and translational activity. This convergence aligns with earlier reports indicating that transcriptomic and proteomic divergence between dormant and germinating seeds is most prominent during early imbibition and tends to attenuate over time (Rajjou *et al*., 2004; Nakabayashi *et al*., 2005). It suggests that the molecular programs dictating dormancy maintenance or release are likely initiated rapidly and rely on early translation events.

GO enrichment analysis of genes consistently downregulated at both transcriptional and translational levels points to a suppression of stress-response pathways and hormone-related signaling, particularly abscisic acid (ABA), which is known to enforce dormancy (Finkelstein *et al*., 2008). In contrast, genes involved in translation, ribosome biogenesis, and protein synthesis were coordinately upregulated, consistent with the preparation of the translational apparatus for active germination (Bai *et al*., 2020; Sano *et al*., 2020). Additionally, the detection of genes involved in RNA processing and translation that are regulated solely at the translational level suggests the action of selective translational control mechanisms. This observation supports the idea that seed germination is not merely driven by *de novo* transcription but also by the differential recruitment of stored mRNAs to ribosomes (Basbouss-Serhal *et al*., 2015). Conversely, transcription-only regulated genes appeared to represent more general cellular functions and were not strongly associated with germination or dormancy, implying that translational control confers specificity to the developmental program.

An outcome of our work is the discovery that many master regulators of germination are under translational control, rather than transcriptional regulation. For example, among them, *GA20ox1, GA3ox1, GA3ox2*, critical actors GA biosynthesis, show negligible translation in dormant seeds but exhibit sharp increases in ribosome occupancy following dormancy release (Fig. 3C). These results extend previous transcriptional studies by showing that the production of germination-promoting proteins is not merely controlled at the transcript level but is selectively gated at the level of ribosome association (Tyler *et al*., 2004; Ariizumi *et al*., 2013). Conversely, transcripts encoding key dormancy factors such as *DOG1* and *ABI5* were found to be highly translated in dry dormant seeds, despite limited environmental cues (Fig. 3B). Our data reveal that translation of these dormancy-enforcing transcripts is sharply decreased following stratification and during imbibition in non-dormant seeds. This dynamic change in ribosome occupancy on mRNAs underscores a decisive reprogramming of translational landscapes during the early phase of seed hydration, consistent with a developmental transition from a dormant to an active state.

### Translational efficiency patterns reflect dormancy status

Our Ribo-seq analysis provides novel insights into the dynamics of translational efficiency (TE) in Arabidopsis seeds during dormancy and germination. TE is a composite parameter that reflects the combined effects of translation initiation, elongation, and termination, and therefore does not report on a single regulatory step. Changes in TE may result from altered initiation rates, but can also arise from differences in ribosome elongation or termination dynamics, for example due to codon usage or ribosome pausing. In addition, TE can be influenced by transcript availability for translation, including regulated sequestration or release of mRNAs from translationally inactive ribonucleoprotein complexes (Bai *et al*., 2020). Here, one of the most striking observations was the globally higher TE in dormant seeds compared to non-dormant seeds, both after stratification and throughout early imbibition (Fig. 4A and 5D). This suggests that dormancy is not simply a state of low metabolic activity, but rather one involving active and selective translation of specific mRNAs. This is in agreement with earlier reports showing that dormant seeds retain high translational potential and maintain active control of protein synthesis through ribosome-associated transcripts and responsive mRNAs (Galland *et al*., 2014; Basbouss-Serhal *et al*., 2015; Bai *et al*., 2020). The ability to maintain translational control may be critical for stress tolerance and for readiness to respond rapidly to favorable germination cues. Intriguingly, dry seeds also displayed high average TE (Cluster 3, Fig. 4A and Cluster 1, Fig. 4B), supporting the concept that seed-stored mRNAs are not passively preserved, but are actively engaged with the translation machinery. This pre-loading of ribosomes in the dry state may serve as a “priming” mechanism, preparing the seed proteome for immediate response upon imbibition, a hypothesis that aligns with the concept of translational poising found in other developmental contexts (Sano *et al*., 2020)

To further explore this, we categorized genes based on ribosome density in dry seeds. Transcripts in the lowest TE decile (1^st^ decile) in dry seeds revealed distinct behaviors (Fig. 4A). A large cluster of these genes showed increased TE specifically during imbibition of dormant seeds, but not in non-dormant ones. GO enrichment of this cluster identified key processes such as translation initiation, ribosome biogenesis, and protein folding (Fig. 4B), functions that are essential for seed rehydration and the eventual transition to active growth. This suggests that dormant seeds selectively enhance translation of components of the protein synthesis machinery, possibly to preserve readiness for germination under strict environmental thresholds. Transcripts from the highest TE group (10^th^ decile) in dry seeds tended to show declining or inconsistent TE during stratification and imbibition, regardless of dormancy status. These genes did not cluster under a specific functional category, as revealed by broad and unstructured GO terms (Fig. 4D). This implies that high TE in dry seeds does not necessarily predict sustained translational activity during imbibition, especially for transcripts not directly involved in dormancy control or early germination.

Taken together, these results support a model in which translational efficiency is a key regulatory layer distinguishing dormant from non-dormant seed states. Importantly, changes in translational efficiency likely reflect multiple underlying mechanisms, including the selective sequestration of mRNAs in translationally inactive ribonucleoprotein complexes, as well as transcript-intrinsic features such as uORFs or secondary structure that modulate translation initiation and elongation. Thus, differential TE does not necessarily imply altered translational capacity per se, but rather highlights the integration of transcript availability and sequence-dependent regulatory controls during seed dormancy and germination. The ability of dormant seeds to selectively increase TE for genes involved in translation and protein processing indicates a tightly regulated post-transcriptional program that may delay germination until appropriate environmental conditions are sensed. In contrast, non-dormant seeds appear to engage in a more general downregulation of TE, possibly reflecting their commitment to germination and proteome restructuring based on *de novo* transcription (Nakabayashi *et al*., 2005; Sano *et al*., 2020).

### Ribosome occupancy in dry seeds reveals a poised translational state and uORFs serve as key translational checkpoints

Remarkably, contrary to expectations, we observed that ribosomes in dry dormant seeds are not idle. Instead, they are strongly enriched at canonical translation initiation sites (the start codons) and within the early coding regions of many mRNAs, forming sharp peaks indicative of active or paused translation complexes (Fig. 5). In dry seeds, our data reveal pronounced P-site enrichment at start codons, supporting a model (Inset Fig. 5A) in which initiating ribosomes are stalled with the initiator Met-tRNAᵢ positioned in the P-site while the A-site remains unoccupied. This suggests that dry seeds, although lacking visible metabolic activity, retain a highly organized translational machinery that is "primed" and ready to resume protein synthesis upon imbibition. This contrasts with the assumption made by (Bai *et al*., 2020), who proposed that most stored mRNAs in dry seeds are associated with monosomes. However, their conclusions were based on polysome profiling, a method that does not provide precise information about ribosome occupancy and position on individual transcripts. Our codon level positioning of ribosomes by Ribo-seq aligns with earlier biochemical work suggesting the presence of preassembled ribosome-mRNA complexes in dry seeds (Galland *et al*., 2014), and provides transcriptome-wide confirmation at nucleotide resolution, reinforcing the model of a poised translational state as a key mechanism of dormancy maintenance and rapid germination response.

A defining feature of translational regulation in our dataset is the widespread presence and active translation of uORFs, which modulate access to main coding regions (von Arnim *et al*., 2014). Mapping of ribosome footprints revealed extensive ribosome occupancy within 5′ UTRs. These signals were enriched around start codons of uORFs and showed conserved phasing, consistent with genuine translation events. Critically, we observed a general dynamic decrease in the ratio of mORF-to-uORF translation during seed imbibition, with a sharp shift at 15 hai, whether weeds were dormant or not (Fig. 6A). Seed dormancy release (stratification) revealed both an increase and a decrease in this ratio for a similar proportion of transcripts (clusters 3, 5, 2 and 1,4 in Fig. 6B, respectively), suggesting a tight control of translation by uORFs during this process. This suggests that dormancy alleviation is not merely the result of transcriptional activation but involves the lifting of uORF-mediated repression, allowing previously inaccessible mRNAs to be efficiently translated. Hierarchical clustering indicated that uORF activity is tightly linked to physiological status and may serve as a checkpoint between dormancy and germination competence.

We functionally validated uORF repression using two regulatory genes (*MARD1* and *PAO4*) each containing conserved AUG-initiated uORFs. Ribo-seq confirmed ribosome occupancy on these uORFs *in vivo*, and luciferase reporter assays in protoplasts showed that mutation of the uORF start codon led to significant decrease of translation, confirming their functional role. *MARD1*, a key ABA-responsive dormancy factor (He & Gan, 2004), showed stronger repression, consistent with its translational silencing during dormancy. *PAO4*, which is involved in redox regulation via polyamine catabolism (Benkő *et al*., 2022), displayed more variable TE, potentially reflecting its role in stress-response tuning during germination. These results place uORFs as molecular regulators controlling the timing of protein production, and their activity as a core determinant in dormancy maintenance and release.

## Conclusion

This study of ribosome profiling analysis offers unique insights into the translational landscape of seed dormancy and germination. Our results reveal that ribosomes in dry Arabidopsis seeds are not inactive, but rather strategically positioned on start codons and coding sequences of key dormancy- and germination-related genes, establishing a primed translational state.

We further show that germination is driven by selective translation, with key regulatory genes, including *GA20ox, GA3ox, DOG 1*, or *ABI5*, under tight translational control. Moreover, we demonstrate that uORF-mediated repression is a major mechanism for translational gating, dynamically modulated during seed imbibition and stratification.

Taken together, our findings redefine the role of translation in seed biology, highlighting it as a central and dynamic regulatory layer. By uncovering the interplay between ribosome positioning, translational efficiency, and uORF repression, we establish a mechanistic framework that explains how seeds remain dormant yet poised for activation, and how translation orchestrates the transition to germination.

These insights have broad implications for crop management, seed technology, and fundamental plant developmental biology. Targeting translational regulators or modifying uORF motifs may offer new strategies for engineering seed behavior under variable environmental conditions.

## Acknowledgements

J.B. was funded by Marie Curie European Economic Community Fellowship PIOF-GA-2012-327954 and US National Science Foundation MCB-1021969.

## Notes

### Competing Interest Statement

The authors have declared no competing interest.

