## Supplemental Information for "Ribosome profiling reveals distinct translational programs underlying Arabidopsis seed dormancy and germination"

### Supplemental Figures and Tables

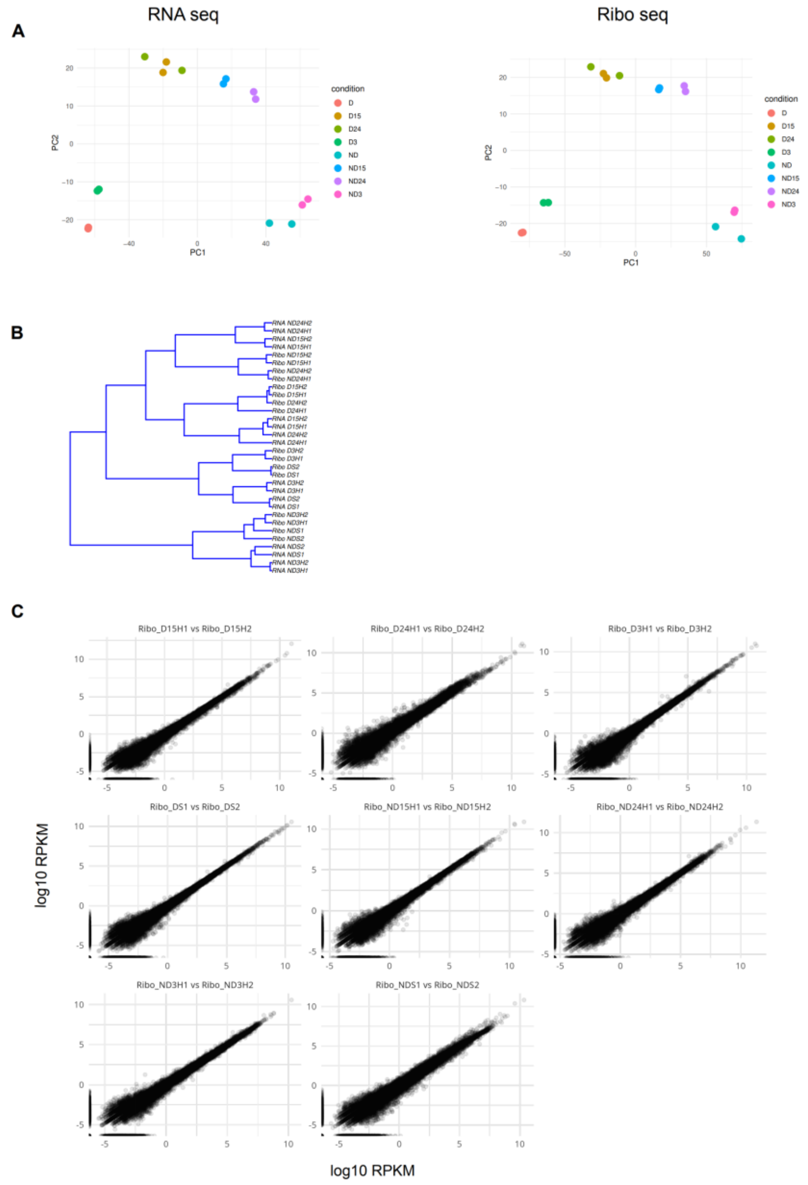


**Figure S1.**

A) PCA analysis of the Ribo seq (Ribo) and RNA-seq (RNA) samples at 0, 3, 15 and 24 hours after imbibition (hai)

B) Correlation tree of the RNA and Ribo-seq samples

C) Reproducibility of the Ribo-seq replicates

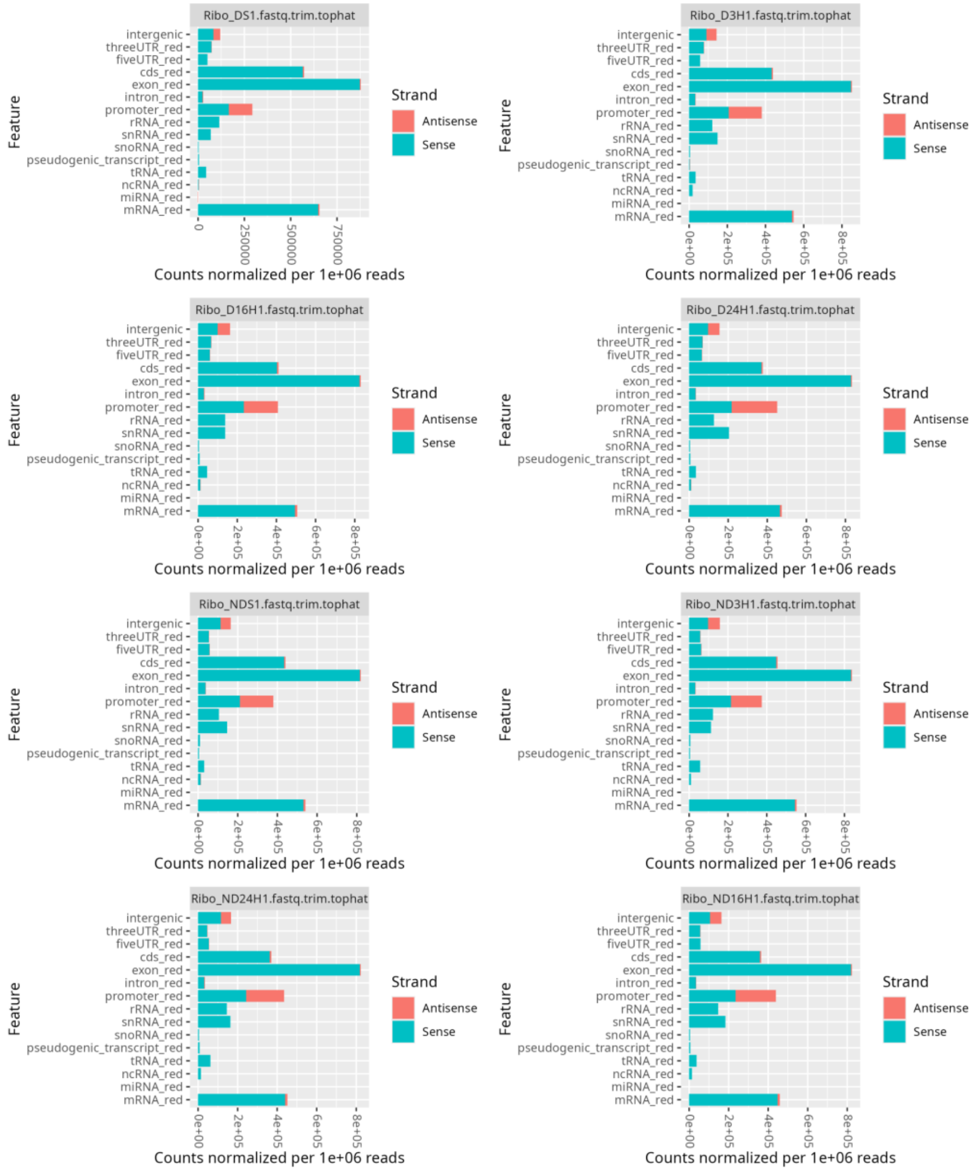


**Figure S2**
Counts of Ribo-seq reads mapping on different genome features

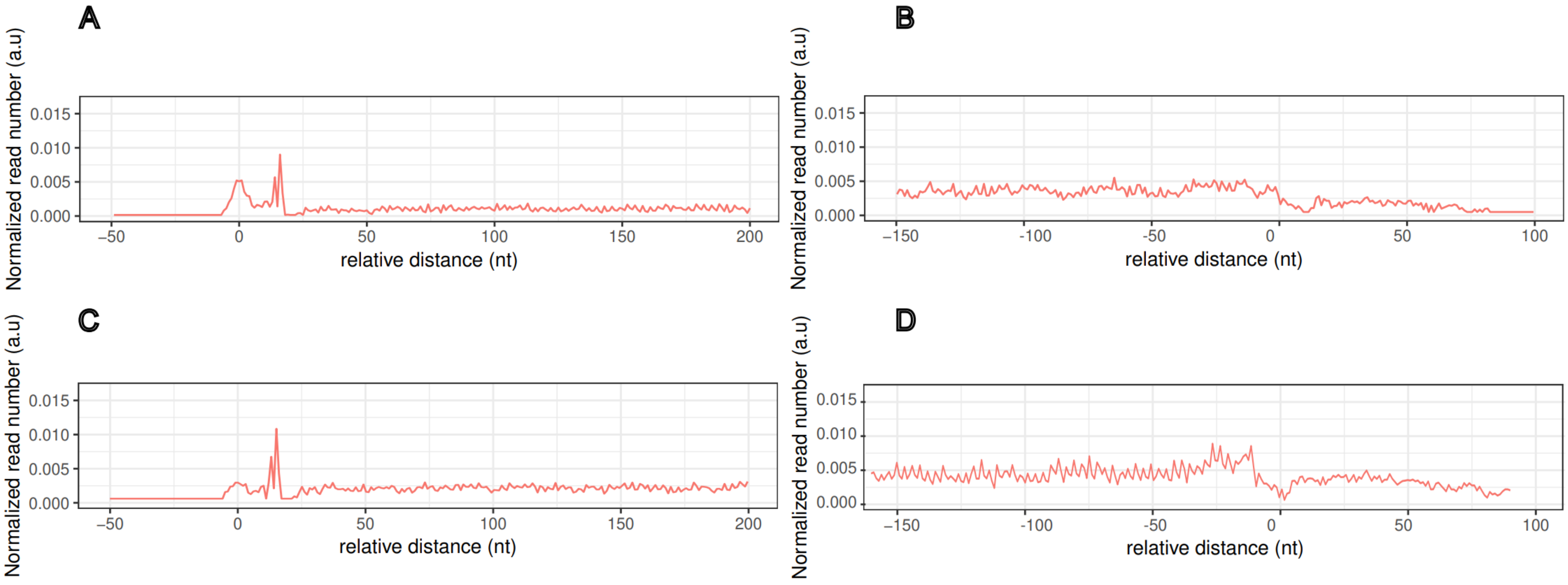


**Figure S3.** Metagene analysis of P-Sites distribution around start (**A, C**) and stop (**B, D**) codon on genes with low TE (1st decile; **A, B**) and high TE (10th decile; **C,D**) in dry seeds

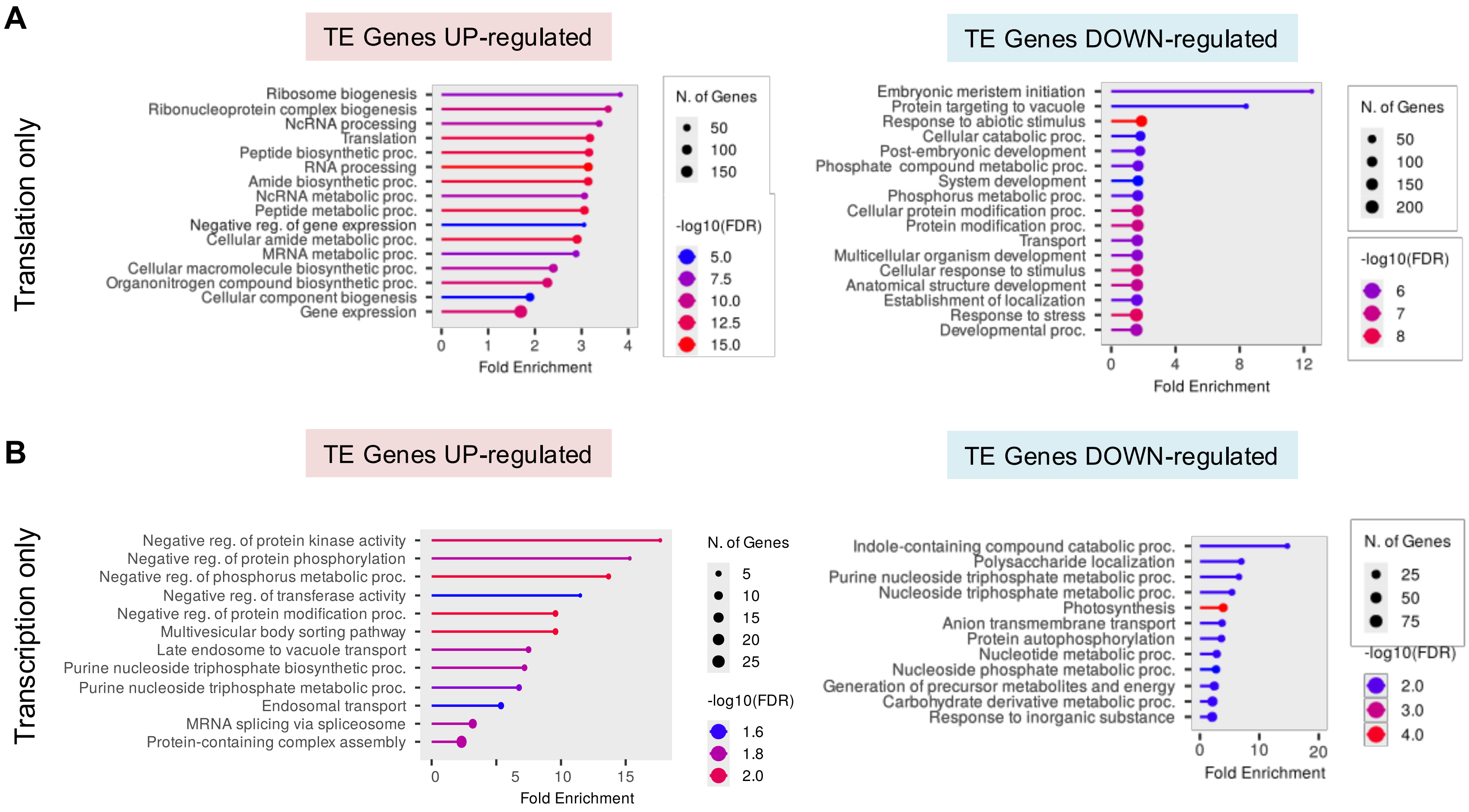


**Figure S4. A)** GO analysis of genes only translationally downregulated and upregulated during stratification (all sample times combined, dry, 3H, 15H and 24H). **B)** GO analysis of genes only transcriptionally downregulated and upregulated during stratification (all sample times combined, dry, 3H, 15H and 24H).

**Table S1. Primers used in this study.**

| Name | Sequence (5′ to 3′) |
|---|---|
| 5'UTR_For_PAO1_uORF | GTACTAGTgtcaccaaacccttcatcttcttc |
| 5'UTR_Rev_PAO1_uORF | TGCCATGGgtgaagattttgttggaaagaaacag |
| 5'UTR_For_MARD1_uORF | GTACTAGTctcgctctttggaaggaataaag |
| 5'UTR_Rev_MARD1_uORF | gGCGGCCGCagttaaagaccggtgagggtaa |
| 5'UTR_For_PAO1_mut-uORF | CGTGATCGGAGCCCTTTTTTTTTTG |
| 5'UTR_Rev_PAO1_mut-uORF | GAATTGTAAGATCTCCGCCGCGAACG |
| 5'UTR_For_MARD1_mut-uORF | CTTTCCTCTCTCCAAAGAAGCTACTGAG |
| 5'UTR_Rev_MARD1_mut-uORF | CAATGAGTCTGCGTAAGAGAAAATATC |
| Seq Luciferase | CGCAACTGCAACTCCGATAAA |
